# Single *JAK2*-V617F hematopoietic stem cells can initiate MPN in transplantations into non-conditioned recipient mice

**DOI:** 10.1101/2025.06.08.657469

**Authors:** Quentin Kimmerlin, Morgane Hilpert, Nils Hansen, Alexandre Guy, Marc Usart, Jan Stetka, Patryk Sobieralski, Tiago Almeida Fonseca, Hui Hao-Shen, Radek C. Skoda

## Abstract

Myeloproliferative neoplasms (MPN) are clonal disorders of hematopoietic stem cells (HSC) that are most frequently caused by acquired somatic mutations in *JAK2*. A number of conditional mouse models of *JAK2-*V617F-driven MPN have been generated that rely on Cre-LoxP mediated activation, resulting in polyclonal disease. To more closely mimic the monoclonal origin of human MPN, transplantations of single purified *JAK2*-mutant HSCs or bone marrow (BM) at limiting dilutions into lethally irradiated recipient mice have been previously performed. However, irradiation is known to alter the BM microenvironment and also to induce transient aplasia accompanied by elevated cytokine levels that promotes the expansion of the mutant clone. To overcome these limitations, we examined whether *JAK2-*V617F-mutant HSCs are able to engraft and initiate MPN in non-conditioned recipients. We found that BM from two different MPN models, one expressing the human *JAK2*-V617F, and another expressing the mouse *Jak2*-V617F, efficiently engrafted and initiated MPN in non-irradiated immunocompromised *Rag2^−/−^* recipients. MPN evolved even in transplantations at limiting dilutions, showing high competitiveness of single *JAK2*-mutant HSCs. Thus, *JAK2*-V617F mutant HSCs can outcompete resident non-mutated HSCs in the absence of elevated cytokine levels and without the need of emptying stem cell niches by irradiation. However, only BM from mice expressing the mouse *Jak2*-V617F engrafted and initiated disease in non-conditioned C57BL/6 mice, while BM from mice expressing the human *JAK2*-V617F was rejected, indicating that mouse *Jak2*-V617F is ignored by the immune surveillance. These results provide a possible explanation why *JAK2*-V617F is so frequently found in healthy individuals with clonal hematopoiesis.

## INTRODUCTION

Myeloproliferative neoplasms (MPN) are clonal disorders of hematopoietic stem cells (HSC) that are caused by acquired somatic mutations in *JAK2*, *MPL*, or *CALR* genes, which result in hyperproliferation of erythroid, megakaryocytic, or myeloid lineages.^1,2^ The *JAK2-* V617F substitution is the most frequent acquired driver mutation, found in about 70% of all MPN patients.^3–6^ This gain-of-function mutation leads to cytokine hypersensitivity, manifesting in three distinct MPN phenotypes: polycythemia vera (PV), where increased erythrocyte production predominates, essential thrombocythemia (ET), marked by excessive platelet production, and primary myelofibrosis (PMF) with extramedullary hematopoiesis.^7,8^

A number of mouse models of *JAK2-*V617F driven MPN have been generated (reviewed in refs.^9–14^). The majority of these models rely on conditional alleles that need to be activated by the Cre-LoxP recombinase system resulting in polyclonal disease, because *JAK2*-V617F is simultaneously induced in a large number of individual HSCs. We previously established monoclonal models of the disease by transplanting single purified *JAK2*-mutant HSCs or bone marrow (BM) at limiting dilutions into lethally irradiated recipient mice.^15^ However, irradiation is known to disrupt the BM microenvironment and also to induce transient aplasia accompanied by elevated cytokine levels, which may artificially promote the expansion of the transplanted *JAK2*-mutant cells.

To overcome these limitations, we sought to establish MPN models originating from single HSCs by transplantation of *JAK2*-mutant cells into non-conditioned recipient mice. As donors, we used two mouse lines, one expressing the human *JAK2*-V617F,^16^ and another expressing the mouse *Jak2*-V617F.^17^ As recipients, we used non-irradiated immunocompromised *Rag2^−/−^* mice,^18^ or fully immunocompetent C57BL/6 mice. Our findings indicate that *JAK2*-V617F HSCs can outcompete resident *JAK2* wildtype hematopoiesis without pre-conditioning of the host.

## MATERIAL AND METHODS

### Mice

We used mice with conditional *JAK2*-V617F “flip-flop” allele (*FF1*),^16^ or conditional *Jak2*-V617F knock-in (*K*i) allele.^17^ These mice were crossed with the tamoxifen-inducible *SclCre^ER^* mice,^19^ the *VavCre* mice,^20^or the interferon-inducible *MxCre* mice.^21^ *FF1* and *Ki* mice were also crossed with the *UBC-GFP* strain that constitutively express the green fluorescent protein (GFP) in all hematopoietic lineages^22^, allowing us to track donor-derived chimerism in bone marrow transplantation studies. *MxCre* mice, *SclCre^ER^*mice and *UBC-GFP* mice have been backcrossed into the C57BL/6 background for 12 generations and the purity of the C57BL/6 background was verified using a 1449 SNP Illumina BeadChip panel and the SNaP-Map Software at the DartMouse Speed Congenic Core Facility at The Geisel School of Medicine at Dartmouth. Cre^ER^ enzymatic activity was induced by intraperitoneal injection of 2 mg tamoxifen (Sigma Aldrich) for 5 consecutive days and Cre expression in *MxCre* mutant mice was induced by intraperitoneal injection of 300µg polyinosine-polycytosine (pIpC) 3 times every second day. C57BL/6 mice (8 to 10 weeks old) were purchased from Janvier Labs, Saint Berthevin, France, or were obtained from in house breeding. *Rag2*^−/−^ mice on C57BL/6 background were from Taconic.^18^ All mice used in this study were maintained under specific pathogen-free conditions and in accordance with Swiss federal regulations. All animal experiments were approved by the Cantonal Veterinary Office of Basel-Stadt, Switzerland.

### Bone marrow transplantations

Bone marrow cells were harvested from donor mice 8-12 weeks after induction by tamoxifen or pIpC by crushing long bones (two tibias, two femurs) and the pelvis using a mortar and pestle in staining medium (PBS + 5% FCS). The cell suspension was filtered through a 70 µm nylon mesh to obtain single cells, followed by red blood cell lysis with ACK lysing buffer (Sigma-Aldrich, St. Louis, MO, USA) for 15 minutes and subsequent washing. The cell suspension was used directly for transplantation or FACS-sorted to isolate HSCs (Lin− C-kit+ Sca1+ CD48− CD150+). Donor cells were finally resuspended in cold Phosphate-Buffered Saline (PBS) (Sigma-Aldrich) and injected intravenously into recipient mice via the tail vein (the number of cells injected is specified in each figure). Blood samples were collected every four weeks to monitor blood counts and chimerism. Secondary transplantations were performed using bone marrow cells harvested from primary recipients at termination. Engraftment was defined as GFP-chimerism in Gr1+ cells >5% at 16 weeks or as *JAK2-*V617F chimerism in total peripheral blood leukocyte DNA >2%. MPN phenotype in mice that engrafted was defined as hemoglobin >170 g/L, or platelet counts >2.2 × 10^9^/L, or neutrophils >2.5 × 10^9^/L at any time point.

### Blood collection and analysis

EDTA-anticoagulated blood was collected by making a small incision in the lateral tail vein of restrained mice. Blood samples were analyzed for complete blood counts using an Advia 120 Hematology Analyzer using the Multispecies Software (Bayer, Leverkusen, Germany) or a Sysmex XN-1000™ Automated Hematology Analyzer (Sysmex Corporation, Kobe, Japan) and analyzed for chimerism by flow cytometry.

### Flow cytometry

To isolate LT-HSCs (defined as Lin− C-kit+ Sca1+ CD48− CD150+), bone marrow cells were first incubated with biotinylated Lineage-negative antibody cocktail consisting of anti-CD3e (clone 17-A2), anti-CD4 (clone L3T4), anti-CD8 (clone 53-6.72), anti-B220 (clone RA3-6B2), anti-TER-119 (clone Ter-119) and anti-Gr-1 (clone RB6-8C5) antibodies (BioLegend, San Diego, CA, USA). The cells were then stained with BV711-conjugated anti-c-Kit (clone 2B8), APC-Cy7-conjugated anti-Sca-1 (clone E13-161.7), AF700-conjugated anti-CD48 (clone HM48-1), and PE-conjugated anti-CD150 (clone TC15-12F12.2) antibodies (BioLegend). This was followed by incubation with Pacific Blue-conjugated streptavidin (Thermo Fisher Scientific, Waltham, MA, USA), after which the cells were washed, resuspended in Sytox Blue (Thermo Fisher Scientific), and sorted on a FACSAria III cell sorter (BD Biosciences, Franklin Lakes, NJ, USA).

To analyze peripheral blood chimerism, EDTA-anticoagulated blood was stained with APC-coupled anti-Ter119 (clone Ter-119) and PE-coupled anti-CD61 (clone 2C9.G2) antibodies or with Pe-Cy7-coupled anti-Gr1 (clone RB6-8C5), APC-coupled anti-CD11b (clone M1/70) antibodies (BioLegend). The cells were washed, resuspended in Sytox Blue (Thermo Fisher Scientific), and recorded on a LSRFortessa Flow Cytometer (BD Biosciences).

To analyze LT-HSC (defined as Lin− C-kit+ Sca1+ CD48− CD150+) and LSK (defined as Lin− C-kit+ Sca1+) cells after termination, bone marrow and spleen cell suspensions were first incubated with biotinylated Lineage-negative antibody cocktail consisting of anti-CD3e (clone 17-A2), anti-CD4 (clone L3T4), anti-CD8 (clone 53-6.72), anti-B220 (clone RA3-6B2), anti-TER119 (clone Ter-119) and anti-Gr-1 (clone RB6-8C5) antibodies (BioLegend). The cells were subsequently stained with BV711-coupled anti-c-Kit (clone 2B8), APC-Cy7-coupled anti Sca-1 (clone E13-161.7), AF700-coupled anti-CD48 (clone HM48-1), PE-Cy7-coupled anti-CD150 (clone TC15-12F12.2) antibodies (BioLegend). This was followed by incubation with Pacific Blue-conjugated streptavidin (Thermo Fisher Scientific), after which the cells were washed, resuspended in Sytox Blue (Thermo Fisher Scientific), and recorded on a LSRFortessa Flow Cytometer (BD Biosciences) or on an Aurora Spectral Flow Cytometer (Cytek Biosciences, Fremont, CA, USA).

To analyze erythroid progenitors (EryP, CD71+Ter119+), bone marrow and spleen cell suspensions were first incubated with APC-coupled anti-Ter119 (clone Ter-119) and PE-coupled anti-CD71 (clone RI7217) antibodies (BioLegend), then washed, resuspended in Sytox Blue (Thermo Fisher Scientific), and recorded on a LSRFortessa Flow Cytometer (BD Biosciences) or on an Aurora Spectral Flow Cytometer (Cytek Biosciences, Fremont, CA, USA). All data were analyzed using FlowJo (version 10.0.08) software (Treestar, Ashland, OR, USA).

### *JAK2*-V617F chimerism in the absence of GFP

To assess *JAK2-*V617F (*JAK2*-VF) chimerism in the blood, bone marrow, and spleen of non-conditioned mice transplanted with *SclCre;FF1* BM donor cells (heterozygous for the human *JAK2-*V617F allele), we first established a standard curve using quantitative PCR (qPCR). PCR amplification was performed using primers specific for the *FF1* allele (5′-TCA CCA ACA TTA CAG AGG CCT ACT C-3′ and 5′-GCC AAG GCT TTC ATT AAA TAT CAA A-3′) and the housekeeping mouse gene *Gusb* (5′-ATA AGA CGC ATC AGA AGC CG-3′ and 5′-ACT CCT CAC TGA ACA TGC GA-3′). To generate the standard curve, genomic DNA was extracted from whole blood of *WT* and induced *SclCre*;*FF1* mice (DNeasy Blood & Tissue Kit, Qiagen) and mixed in defined ratios. qPCR was performed on an ABI 7500 analyzer (Applied Biosystems), and the relative quantity of the FF1 allele (ΔΔCt method) was plotted against the predefined *FF1/WT* DNA ratios to establish the standard curve. Peripheral blood, bone marrow, and spleen DNA from transplanted mice were processed similarly, and chimerism levels were determined by comparing the relative *FF1* quantity to the standard curve.

To assess *Jak2*-V617F (*Jak2*-VF) chimerism in the blood, bone marrow and spleen of non-conditioned mice transplanted with *SclCre;Ki* BM donor cells (heterozygous for the mouse *Jak2-*V617F allele), we used the size difference of the intron between exon 13 (mutation site) and 14, which is slightly longer for the *Ki* allele compared to *WT*. PCR amplification using primers flanking this region (5’-TGT CTT ACT AAA GCC CAG GTG ATG G-3’ and 5’-GCT CCA GGG TTA CAC GAG TC-3’),^17^ allowed us to generate a standard curve to quantify JAK2VF chimerism. For this, genomic DNA was extracted from whole blood of *WT* and induced *SclCre*;*Ki* mice (DNeasy Blood & Tissue Kit, Qiagen) and mixed in defined ratios. PCR was then performed, and the resulting products were separated by capillary electrophoresis (Agilent 4200 TapeStation System). Band intensities were quantified, and the *Ki/WT* intensity ratio was plotted against the predefined *Ki/WT* DNA ratios to establish the standard curve. Peripheral blood, bone marrow, and spleen DNA from transplanted mice were processed similarly, and chimerism levels were determined by comparing their *Ki/WT* band intensity ratio to the standard curve.

### Statistical analysis

Statistical comparisons were performed as indicated in the figure legends. Data were analyzed and plotted using Prism software version 10.4.1 (GraphPad Inc.). All data are represented as mean ± SEM. Significance is denoted with asterisks (*P<0.05, **P<0.01, ***P < 0.001, ****P < 0.0001).

## RESULTS

### Bone marrow transplantation into non-conditioned Rag2^−/−^ recipient mice

To test whether the *JAK2*-mutant HSC can engraft and initiate MPN in non-conditioned hosts and to avoid possible rejection, we selected non-irradiated *Rag2^−/−^* mice as the recipients because they are immuno-deficient since they lack B and T cells. As donors we used two different MPN models, one expressing the human *JAK2*-V617F (*FF1*),^16^ and another expressing the mouse *Jak2*-V617F (*Ki*).^17^ Total bone marrow (BM) cells harvested from induced *SclCre;FF1;GFP* or *SclCre;Ki;GFP* donors all engrafted and, with the exception of two *SclCre;FF1;GFP* recipients, rapidly produced MPN phenotypes (Figure 1A and 1B). In fact, because 2 *SclCre;FF1;GFP* recipients died early and other *SclCre;FF1;GFP* and *SclCre;Ki;GFP* recipients looked sick, we terminated the experiment already 12 weeks after transplantation. In contrast, recipients of BM from *VavCre;FF1;GFP* and *VavCre;Ki;GFP* donors showed a milder phenotype and were followed for 28 weeks. All *VavCre;FF1;GFP* engrafted and developed a thrombocytosis phenotype, while only 5 of 8 *VavCre;Ki;GFP* recipients developed MPN. Mice with MPN phenotype also displayed increased spleen weights (Figure 1C).

**Figure 1.**
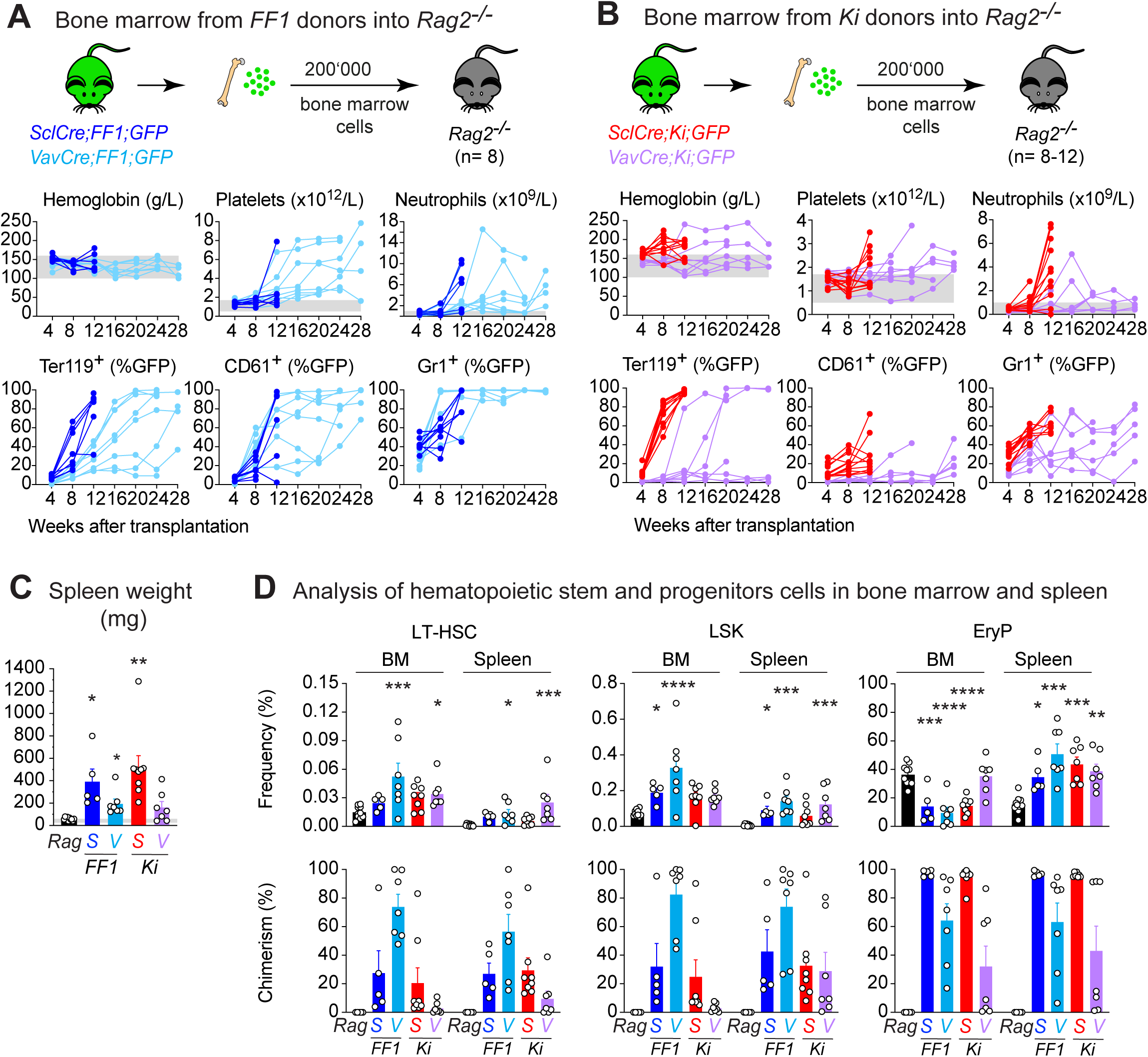
Transplantations of *JAK2*-V617F mutant total bone marrow cells into non-conditioned *Rag2*^−/−^ mice. (**A and B**) Experimental design. A total of 200,000 unfractionated bone marrow (BM) cells obtained from *SclCre;FF1;GFP* (blue), *VavCre;FF1;GFP* (cyan), *SclCre;Ki;GFP* (red) or *VavCre;Ki;GFP* (purple) donors were transplanted into non-conditioned *Rag2^−/−^*recipient mice. Lower panel: Time course of blood counts and GFP chimerism. Each line represents an individual mouse. (**C**) Spleen weight of *Rag2^−/−^*recipient mice. Non-transplanted *Rag2^−/−^* mice (black) were plotted for comparison. (**D**) Frequencies and GFP chimerism of LT-HSCs (Lin− ckit+ Sca1+ CD150+ CD48−), LSKs (Lin− ckit+ Sca1+) and EryP (erythroid progenitors, CD71+ Ter119+). Non-transplanted *Rag2^−/−^* mice (black) were plotted for comparison. Statistical analyses were performed using one-way ANOVA followed by Fisher’s LSD test, with the non-transplanted *Rag2^−/−^*mice serving as the reference for comparisons. *P < .05; **P < .01; ***P < .001; ****P < .0001.

Analysis of bone marrow and spleen in recipients that developed MPN phenotype revealed modest increases in the frequencies of hematopoietic stem and progenitor cells (HSPCs) when compared to non-transplanted Rag2^−/−^ mice and also variable degrees of GFP-chimerism (Figure 1D). Recipients of BM from *VavCreFF1* donors showed overall higher values in LT-HSCs, which is likely due to the longer follow up until terminal analysis compared to recipients of BM from *SclCre* donors. As a control, we transplanted a 10-fold higher number of BM cells from a wildtype donor (*WT;GFP*), but GFP-chimerism remained below 4% (Supplemental Figure S1). We also performed transplantations with BM from *MxCre;FF1;GFP* and *MxCre;Ki;GFP* donors that had been induced with pIpC and obtained similar results (Supplemental Figure S2). *MxCre;FF1;GFP* recipients showed slower kinetics than *SclCre;FF1;GFP* recipients and not all mice developed MPN, while *MxCre;Ki;GFP* recipients showed similar behavior as *SclCre;Ki;GFP* recipients. Overall, BM from both *FF1* and *Ki* donors engrafted in non-irradiated *Rag2^−/−^* recipient mice with some variation in efficiency depending on the different Cre lines used to activate the conditional alleles.

We performed fluorescence-activated cell sorting (FACS) of HSCs from *SclCre;FF1;GFP* and *MxCre;Ki;GFP* donor mice and also observed successful engraftment and MPN disease in *Rag2^−/−^* recipient mice (Figure 2A-D). The frequencies of engraftment were lower for the *FF1* donors compared to *Ki* donors and overall lower than expected for 100 HSC per transplanted mice. It is at present not clear whether this lower efficiency of engraftment is due to suboptimal HSC purification or whether the non-HSC fraction of BM cells is supporting higher efficiency of HSC engraftment in the transplantations of unfractionated BM. However, analysis of BM and spleen in mice that developed MPN showed that the *JAK2*-mutant clone from *FF1* donor expanded and showed high chimerism in the HSC compartment, whereas the *JAK2*-mutant clone from *Ki* donor showed expansion mainly at later stages of differentiation (Figure 2E).

**Figure 2.**
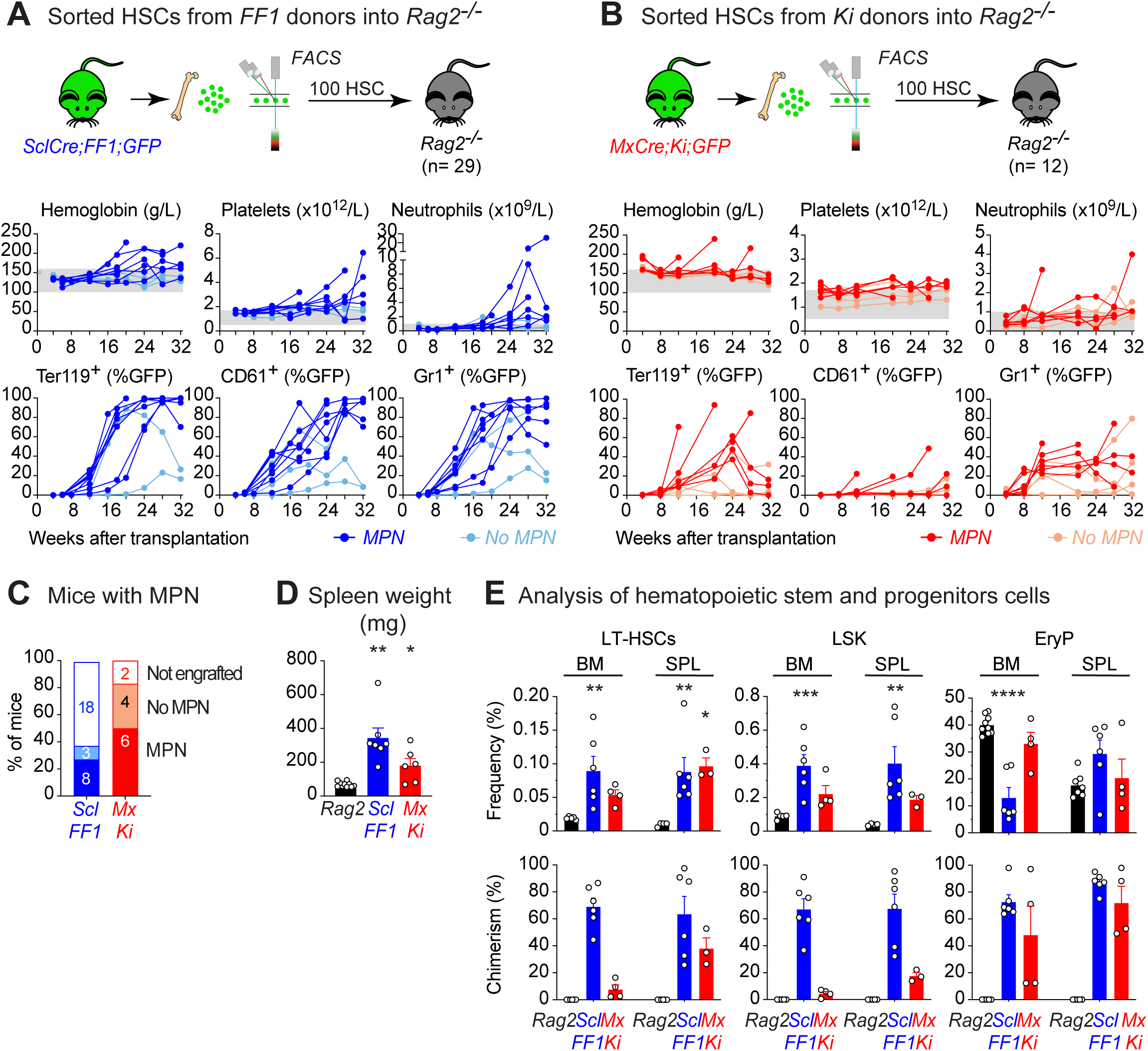
Transplantations of purified LT-HSCs from *JAK2*-V617F donors into non-conditioned *Rag2*^−/−^ mice. **(A and B**) Experimental design. A total of 100 LT-HSCs (Lin− C-kit+ Sca1+ CD48− CD150+) obtained from *SclCre;FF1;GFP* (blue) or *MxCre;Ki;GFP* (red) donors were transplanted into non-conditioned *Rag2^−/−^* recipients. Lower panel: Time course of blood counts and GFP chimerism. Each line represents an individual mouse. Mice that successfully engrafted (GFP chimerism in Gr1+ cells >5% at 16 weeks) and developed MPN are shown in darker colors, whereas those that engrafted without MPN appear in lighter colors. Mice that failed to engraft are not represented. (**C**) Proportions of *Rag2^−/−^*recipients that engrafted and developed MPN are shown in darker colors, while those that engrafted without MPN are in lighter colors. Mice that failed to engraft are represented as unfilled. (**D**) Spleen weight of *Rag2^−/−^*recipient mice that developed MPN phenotype. Non-transplanted *Rag2^−/−^*mice (black) were plotted for comparison. (**E**) Frequencies and GFP chimerism of LT-HSCs, LSKs (Lin− C-kit+ Sca1+) or EryP (erythroid progenitors, CD71+ Ter119+) in recipient mice that developed MPN. Non-transplanted *Rag2^−/−^* mice (black) were plotted for comparison. Statistical analyses were performed using one-way ANOVA followed by Fisher’s LSD test, with the non-transplanted *Rag2^−/−^*mice serving as the reference for comparisons. *P < .05; **P < .01; ***P < .001; ****P < .0001.

### *JAK2-V617F* HSCs at limiting dilutions can initiate MPN in non-conditioned *Rag2*^−/−^ mice

Since MPN in patients originates from a single *JAK2*-mutated HSC, we tested if we can mimic disease initiation in non-conditioned Rag2^−/−^ mice by BM transplantations at limiting dilutions. In previous experiments we found that 8’000 BM cells from *MxCre;FF1;GFP* donors were limiting and contained one or no HSC and engraftment was observed in only 30% of transplanted mice.^15^ With 8’000 BM cells from *SclCre;FF1;GFP* or *SclCre;Ki;GFP* donors injected into non-conditioned *Rag2^−/−^* mice we also observed engraftment and MPN phenotype, albeit at slightly higher frequencies (Figure 3A-C), suggesting that the HSC compartment in the SclCre mice was expanded compared to the MxCre mice. By comparing the frequencies of engraftment (defined as >5% in Gr1+ granulocytes at 16 weeks) between the transplantations with 200’000 BM cells versus 8’000 cells and using the “extreme limiting dilution analysis” algorithm,^23^ we estimated that the numbers of functional HSCs in *SclCre;FF1;GFP* and *SclCre;Ki;GFP* donors were approximately 1/8’000 and 1/4’000 BM cells, respectively. Similar to our previous limiting dilution transplantations in lethally irradiated recipients,^15^ not all Rag2^−/−^ mice that showed engraftment developed MPN (Figure 3C). Mice with MPN phenotype also displayed splenomegaly (Figure 3D), increased frequencies and GFP chimerism of HSPCs (Figure 3E). Interestingly, we observed differences in the expansion of the *JAK2*-mutant clone between *FF1* and *Ki* recipients (Figure 3F). While *Ki* recipients showed expansion of the *JAK2*-mutant (GFP-positive) cells primarily at late stages of erythroid differentiation, most of the *FF1* recipients with erythrocytosis phenotype displayed expansion already in the HSC compartment (right panel). In contrast, *FF1* recipients with thrombocytosis showed less pronounced HSC expansion (left panel). The total numbers of phenotypic LT-HSCs per 10^6^ BM cells was higher in recipients of *FF1* BM than in recipients of Ki BM (Table 1). Because of the large differences in the GFP-chimerism, the actual expansion of the *JAK2*-mutant LT-HSCs clone was highly variable between individual *FF1* recipients (Table 1). Assuming that these mice were transplanted with only 1-2 *JAK2*-mutant HSCs, in some of the recipients, e.g. G4, the LT-HSC must have undergone at least 17-18 cell divisions. Thus, the HSCs from *SclCre;FF1;GFP* donors were able to largely outcompete the non-mutated HSCs in some of the non-conditioned Rag2^−/−^ mice, whereas the HSCs from *SclCre;Ki;GFP* donors were less dominant in the HSC compartment, and instead showed expansion at later stages of differentiation.

**Figure 3.**
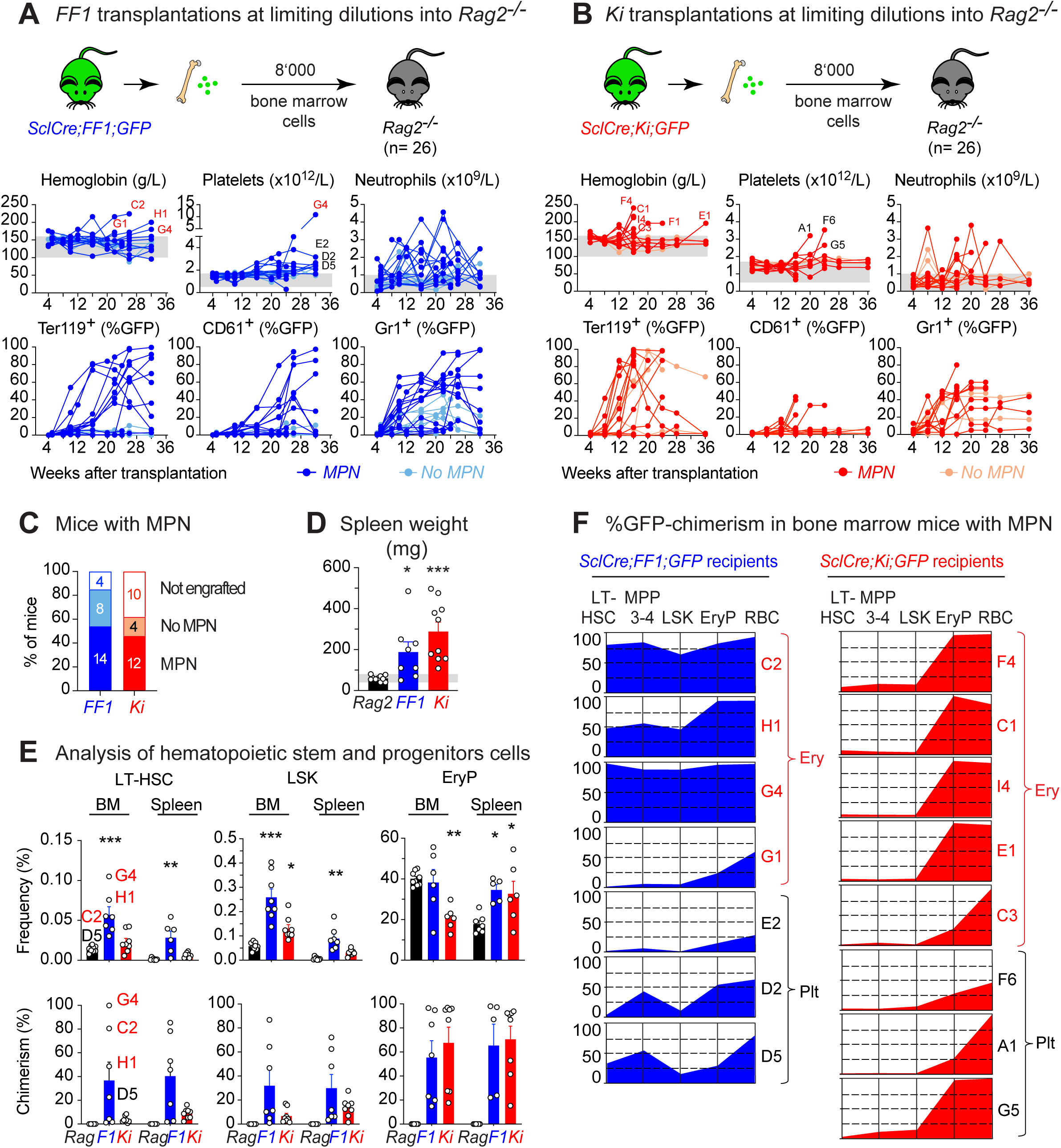
Limiting dilution transplantations of *JAK2*-V617F mutant total bone marrow cells into non-conditioned *Rag2*^−/−^ mice. **(A and B)** Experimental design. A total of 8,000 unfractionated bone marrow (BM) cells obtained from *SclCre;FF1;GFP* (blue) or *SclCre;Ki;GFP* (red) donors were transplanted into non-conditioned *Rag2^−/−^* recipients. Lower panel: Time course of blood counts and GFP chimerism. Each line represents an individual mouse. Mice that successfully engrafted (GFP chimerism in Gr1+ cells >5% at 16 weeks) and developed MPN are shown in darker colors and labeled with a letter and number, whereas those that engrafted without MPN appear in lighter colors. Mice that failed to engraft are not shown. (**C**) Proportions of *Rag2*^□*/*□^ mice that engrafted and developed MPN are shown in darker colors, while those that engrafted without MPN are in lighter colors. Mice that failed to engraft are represented as unfilled. (**D**) Spleen weight of *Rag2^−/−^*recipient mice that developed MPN phenotype. Non-transplanted *Rag2^−/−^*mice (black) were plotted for comparison. (**E**) Frequencies and GFP chimerism of LT-HSCs (Lin− C-kit+ Sca1+ CD48− CD150+), LSKs (Lin− C-kit+ Sca1+) and EryP (erythroid progenitors, CD71+ Ter119+) in recipient mice that developed MPN phenotype. Non-transplanted *Rag2^−/−^* mice (black) were plotted for comparison. (**F**) Analysis of bone marrow from *Rag2^−/−^*recipients that developed an erythrocytosis (Ery) phenotype (identified in red in Figures 2A and 2B) or a phenotype with elevated platelet counts (Plt) (identified in black in Figures 2A and B). GFP-chimerism (%) was assessed in LT-HSC, MPP3/4 (Lin− C-kit+ Sca1+ CD150-), LSK and EryP from bone marrow, and in red blood cells (RBC, Ter119+) from peripheral blood. Statistical analyses were performed using one-way ANOVA followed by Fisher’s LSD test, with the non-transplanted *Rag2^−/−^* mice serving as the reference for comparisons. *P < .05; **P < .01; ***P < .001; ****P < .0001.

**Table 1.**
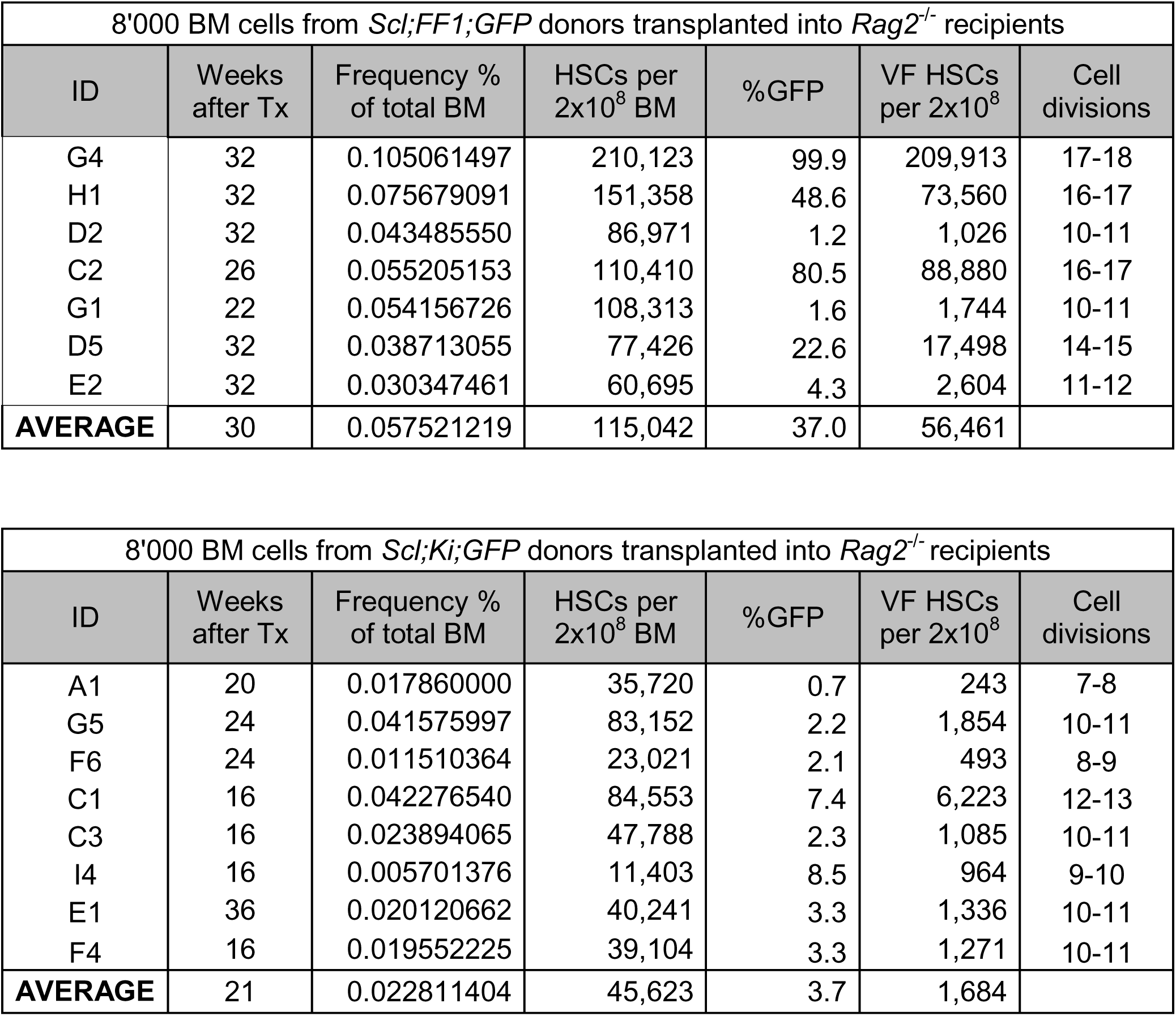
Estimating the expansion of LT-HSCs in limiting dilution transplantations.

| 8'000 BM cells from *Scl;FF1;GFP* donors transplanted into *Rag2<sup>-/-</sup>* recipients |  |  |  |  |  |  |
| --- | --- | --- | --- | --- | --- | --- |
| ID | Weeks after Tx | Frequency % of total BM | HSCs per 2x10<sup>8</sup> BM | %GFP | VF HSCs per 2x10<sup>8</sup> | Cell divisions |
| G4 | 32 | 0.105061497 | 210,123 | 99.9 | 209,913 | 17-18 |
| H1 | 32 | 0.075679091 | 151,358 | 48.6 | 73,560 | 16-17 |
| D2 | 32 | 0.043485550 | 86,971 | 1.2 | 1,026 | 10-11 |
| C2 | 26 | 0.055205153 | 110,410 | 80.5 | 88,880 | 16-17 |
| G1 | 22 | 0.054156726 | 108,313 | 1.6 | 1,744 | 10-11 |
| D5 | 32 | 0.038713055 | 77,426 | 22.6 | 17,498 | 14-15 |
| E2 | 32 | 0.030347461 | 60,695 | 4.3 | 2,604 | 11-12 |
| **AVERAGE** | 30 | 0.057521219 | 115,042 | 37.0 | 56,461 |  |

| 8'000 BM cells from *Scl;Ki;GFP* donors transplanted into *Rag2<sup>-/-</sup>* recipients |  |  |  |  |  |  |
| --- | --- | --- | --- | --- | --- | --- |
| ID | Weeks after Tx | Frequency % of total BM | HSCs per 2x10<sup>8</sup> BM | %GFP | VF HSCs per 2x10<sup>8</sup> | Cell divisions |
| A1 | 20 | 0.017860000 | 35,720 | 0.7 | 243 | 7-8 |
| G5 | 24 | 0.041575997 | 83,152 | 2.2 | 1,854 | 10-11 |
| F6 | 24 | 0.011510364 | 23,021 | 2.1 | 493 | 8-9 |
| C1 | 16 | 0.042276540 | 84,553 | 7.4 | 6,223 | 12-13 |
| C3 | 16 | 0.023894065 | 47,788 | 2.3 | 1,085 | 10-11 |
| I4 | 16 | 0.005701376 | 11,403 | 8.5 | 964 | 9-10 |
| E1 | 36 | 0.020120662 | 40,241 | 3.3 | 1,336 | 10-11 |
| F4 | 16 | 0.019552225 | 39,104 | 3.3 | 1,271 | 10-11 |
| **AVERAGE** | 21 | 0.022811404 | 45,623 | 3.7 | 1,684 |  |

We performed secondary transplantations using three *SclCre;FF1;GFP* donors with MPN phenotype, marked in Figure 3 as G4, H1, and D2. We transplanted 10^6^ BM cells per recipient into non-conditioned *Rag2^−/−^* recipients (Supplemental Figure S3). Based on the frequencies of LT-HSCs in BM of the three donors (Supplemental Figure S3C), we calculated that 10^6^ BM cells from G4 and H1 donors contained ∼1’050 or ∼368 *JAK2*-mutant phenotypic HSCs, respectively, whereas 10^6^ BM cells from the D2 donor contained only ∼5 *JAK2*-mutant HSCs. Recipients of BM from G4 and H1 donors showed engraftment, but only recipients of G4 BM developed MPN phenotype with splenomegaly and high chimerism in BM and spleen (Supplemental Figure S3D-F). In contrast, recipients of BM from donor D2 showed no engraftment and no MPN phenotype (Supplemental Figure S3D-F). Overall, these results suggest that the *JAK2*-mutant HSCs after engrafting and expanding in primary recipients still possess repopulation capacity in secondary recipients, provided that their expansion was sufficient to be present in the 10^6^ BM cells used for the secondary graft.

### *Jak2*-V617F HSCs can initiate MPN also in wildtype C57BL/6 mice

Since the *JAK2*-mutant clone showed high competitive advantage over non-mutant HSC in *Rag2^−/−^* mice, we examined whether transplantation into non-conditioned fully immunocompetent C57BL/6 mice would also be possible. To avoid rejection directed against the Cre-ER fusion protein and/or the GFP protein, we used *MxCre;FF1* and *MxCre;Ki* mice as donors. In these mice the Cre protein is only transiently expressed from the interferon-inducible Mx promoter after induction with pIpC, but is expected to be silent when BM for transplantation is harvested 10 weeks after induction. BM from *MxCre;FF1* donors, which express the human *JAK2*-V617F, did not engraft in C57BL/6 recipients (Figure 4A and C), suggesting that the human JAK2 protein is not tolerated. In contrast, BM from *MxCre;Ki* donors, which express the mouse *Jak2*-V617F, engrafted and in the majority of recipients developed MPN with high hemoglobin, splenomegaly and high *Jak2*-V617F chimerism in BM and spleen (Figure 4B-E). Thus, the fully immunocompetent C57BL/6 mice appear to recognize and reject the human JAK2 protein, whereas the mouse Jak2-V617F protein that differs from WT Jak2 in only one amino acid is ignored by the immune system.

**Figure 4.**
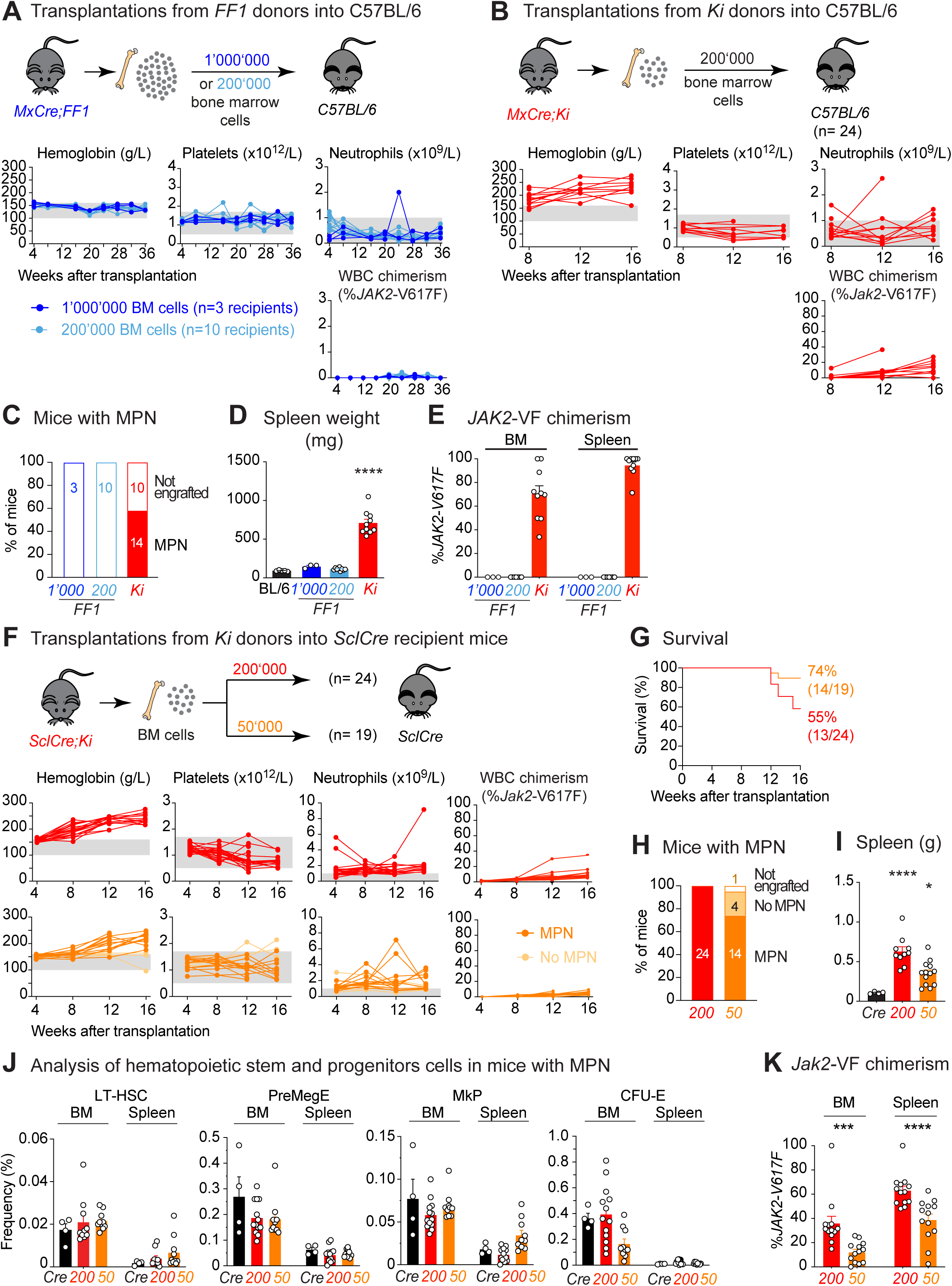
Transplantations of *JAK2*-V617F mutant total bone marrow cells into immunocompetent C57BL/6 mice. **(A and B**) Experimental design. A total of 1’000’000 or 200,000 unfractionated bone marrow (BM) cells obtained from *MxCre;FF1* (blue) or 200,000 BM cells from *MxCre;Ki* (red) donors were transplanted into non-conditioned C57BL/6 recipient mice. Lower panel: Time course of blood counts and *JAK2-*V617F chimerism in white blood cells (WBC). Each line represents an individual mouse. Mice transplanted with *MxCre;FF1* BM cells all failed to engraft and are shown in blue. Mice transplanted with *MxCre;Ki* BM cells that engrafted (*Jak2-*V617F chimerism in WBC DNA >2%) all developed MPN phenotype and are shown in red, while those that failed to engraft are not represented. (**C**) Proportions of C57BL/6 recipient mice that successfully engrafted and developed MPN are shown in darker colors, while those that failed to engraft are shown as unfilled. (**D**) Spleen weights of C57BL/6 recipient mice transplanted with either *MxCre;FF1* BM (which failed to engraft) or *MxCre;Ki* BM that developed MPN. Non-transplanted C57BL/6 mice (black) were plotted for comparison. (**E**) *JAK2*-V617F chimerism was measured by PCR in total BM and spleen DNA from C57BL/6 recipients transplanted with either *MxCre;FF1* BM (which failed to engraft) or *MxCre;Ki* BM that developed MPN. (**F**) Experimental design. A total of 200,000 (red) or 50’000 (orange) unfractionated BM cells obtained from *SclCre;Ki* donors were transplanted into non-conditioned *SclCre* recipient mice. Lower panel: Time course of blood counts and *Jak2*-V617F chimerism in WBC. Each line represents an individual mouse. Mice that engrafted and developed MPN are shown in darker colors, while those that engrafted without MPN are in lighter colors. Mice that failed to engraft are not shown. (**G**) Kaplan-Meier survival plot of transplanted mice. (**H**) Proportions of *SclCre* recipients that engrafted and developed MPN are represented in darker colors, while those that engrafted without MPN are in lighter colors. Mice that failed to engraft are shown as unfilled. (**I**) Spleen weight of *SclCre* recipient mice that developed MPN. Non-transplanted *SclCre* mice (black) were plotted for comparison. (**J**) Frequencies of LT-HSC (Lin− C-kit+ Sca1+ CD48− CD150+), PreMegE (Lin− C-kit+ Sca-1-CD41− CD16/32-CD105− CD150+), MkP (Lin− C-kit+ Sca-1-CD41+ CD150+) and CFU-E (Lin− C-kit+ Sca-1-CD41-CD16/32-CD105+ CD150-) in recipients that developed MPN. Non-transplanted *SclCre* mice (black) were plotted for comparison. (**K**) *Jak2*-V617F chimerism was measured by PCR in total BM and spleen DNA from *SclCre* recipients that developed MPN. Statistical analyses were performed using one-way ANOVA followed by Fisher’s LSD test, with the non-transplanted C57BL6 or *SclCre* mice serving as the reference for comparisons. *P < .05; **P < .01; ***P < .001; ****P < .0001.

After this initial experiment we experienced problems with leakiness of the *MxCre* system, i.e. *MxCre;Ki* already displayed MPN before induction with pIpC. Leakiness of the *MxCre* system is a known problem likely related to activation of the *Mx* promoted by endogenous interferon produced in response to changes in the microbial status of the mouse colony.^21,24^ BM from these pre-induced mice no longer engrafted in C57BL/6 mice, possibly also because continuous expression of Cre as a foreign antigen. We therefore resorted to the *SclCre* system, which shows no leakiness in absence of tamoxifen induction. In order to avoid rejection due to expression of the Cre-ER fusion protein, which is constitutively expressed in hematopoietic cells by the *Scl* promoter, we used *SclCre* mice as the recipients of BM from *SclCre;Ki* mice (Figure 4F). Using 200’000 or 50’000 BM cells per transplant, we observed high frequency of engraftment and MPN phenotype with splenomegaly and shortened survival (Figure 4F-I). Interestingly, in contrast to transplantations into *Rag2^−/−^* mice, the C57BL/6 and *SclCre* recipient mice displayed MPN phenotype already with low *Jak2*-V617F chimerism in total leukocyte DNA (Figure 4B and 4F), as is also the case in some patients with MPN. Analysis of HSPCs in BM and spleen of *SclCre* recipients showed frequencies that were largely unchanged (Figure 4J), but the *Jak2*-V617F chimerism in unfractionated total BM and spleen cells was lower in mice transplanted with 50’000 BM cells compared to 200’000 BM cells (Figure 4K). These experiments show that HSCs that express mouse *Jak2*-V617F are highly competitive in non-conditioned transplantations and escape immuno-surveillance in fully competent wildtype C57BL/6 or *SclCre* recipients, whereas human *JAK2*-V617F expressing cells appear to be recognized and rejected (Figure 5).

**Figure 5.**
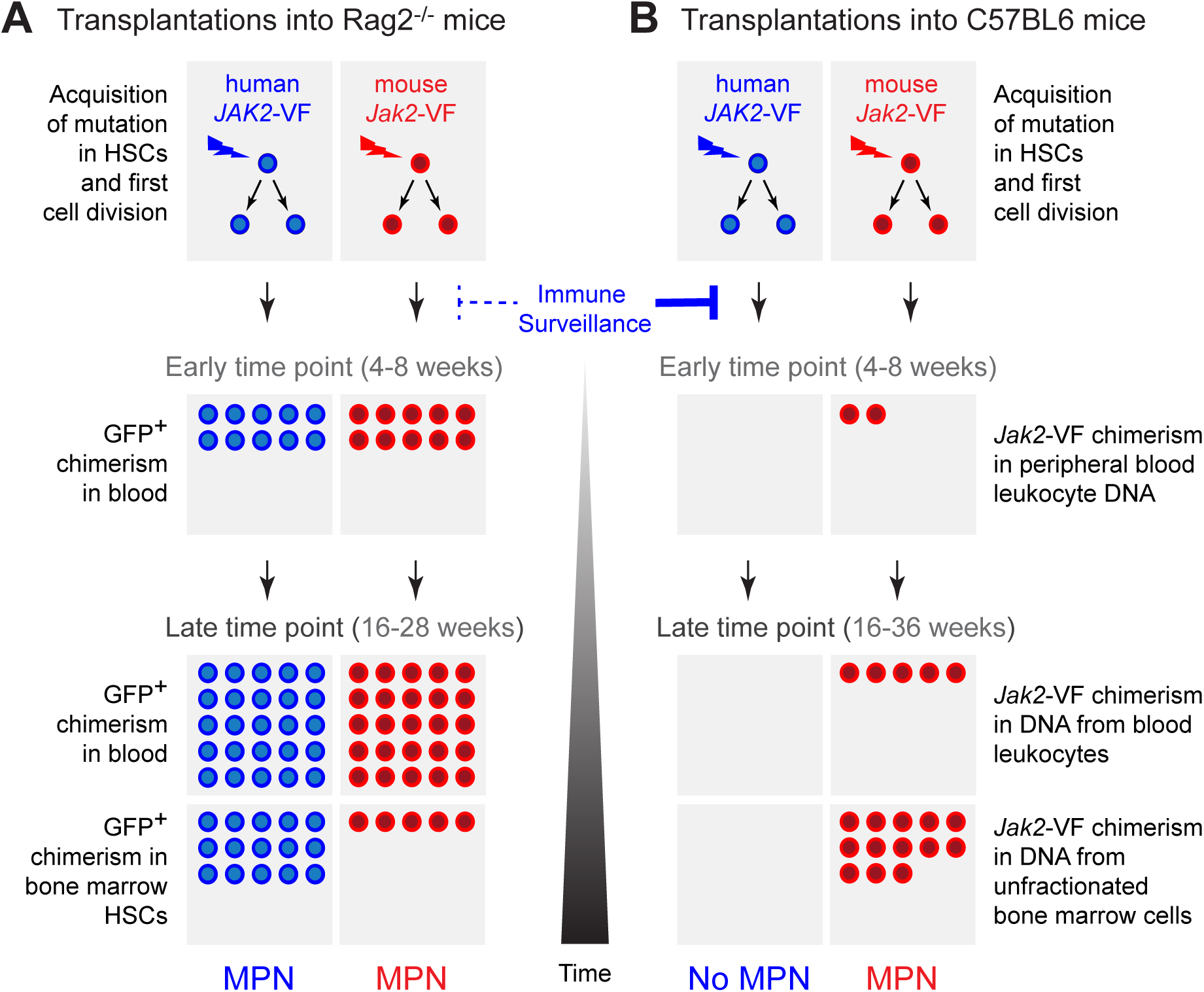
Model summarizing the results obtained in transplantations into non-conditioned Rag2−/− and C57BL/6 mice. (**A**) In *Rag2^−/−^* mice, limiting dilutions of bone marrow (BM) containing a single hematopoietic stem cell (HSC) from *FF1* mice (expressing human *JAK2*-V617F) and from *Ki* mice (expressing mouse *Jak2*-V617F) were sufficient to engraft, expand and cause MPN phenotype over time. HSCs from *FF1* mice expanded well in *Rag2^−/−^* recipients, while HSCs from *Ki* mice showed low chimerism, but strong expansion at later stages of hematopoiesis. (**B**) In non-conditioned C57BL/6 mice, BM cells from *FF1* mice were rejected, indicating that the immune surveillance reacted against cells expressing the xenogeneic human JAK2 protein. In contrast, BM from *Ki* mice expressing the mouse *Jak2*-V617F evaded immune surveillance, engrafted, expanded and the recipient mice developed MPN.

## DISCUSSION

Transplantations of BM from two different MPN mouse models into non-conditioned immunocompromised *Rag2*^−/−^ recipients showed highly efficient engraftment and MPN disease initiation by *JAK2*-mutant cells (Figure 1). This shows that *Rag2*^−/−^ mice that lack B and T cells, exhibited tolerance to both human JAK2 protein and the GFP reporter protein, which enabled monitoring of the lineage contribution of the transplanted JAK2-mutant cells in the recipient mice. The kinetics of MPN disease acquisition in the recipient mice was faster when the conditional *FF1* and *Ki* alleles were activated by tamoxifen in adult *SclCre* donors, compared to recipients that received BM from *VavCre* donor mice (Figure 1A-B), in which the conditional *FF1* and *Ki* alleles are already activated during embryogenesis. This difference in kinetics is likely reflecting the fact that the HSCs in the *VavCre* mice already went through many cell divisions and were already partially exhausted at the time when they were harvested for transplantation. Consistently, a previous report also showed lower efficiency of the *Ki* allele when *VavCre;Ki* donor mice were used in competitive transplantations into lethally irradiated recipients.^17^ Engraftment and MPN disease initiation was also reproduced with sorted HSCs from *FF1* and *Ki* donors (Figure 2). The fact that transplanting 100 purified LT-HSCs showed lower than expected frequencies of engraftment suggests that not all phenotypic LT-HSCs were also functional. This observation is also in line with earlier reports that showed higher efficiency of repopulation when unfractionated BM was used compared to purified LT-HSCs,^25^ possibly also because BM contains cells that may facilitate the engraftment of LT-HSCs.^26^

Although competing with a large excess of non-mutated resident HSCs, transplantations of 8’000 BM cells, i.e. at limiting dilutions, resulted in long-term engraftment and MPN disease initiation in *Rag2*^−/−^ recipient mice (Figure 3). HSCs from *FF1* donors in some recipients massively expanded in numbers (Table 1), while HSCs from *Ki* donors were consistently less dominant in the HSC compartment and these mice showed late expansion mainly at later stages in the erythroid lineage (Figure 3). The basis for the variable expansion of the LT-HSCs from FF1 donors is unclear, but may reflect heterogeneity in the HSC pool.

As previously observed in transplantations of single *JAK2*-V617F mutant HSCs into lethally irradiated recipients,^15^ we again found that at limiting dilutions the recipient Rag2^−/−^ mice showed either erythrocytosis or thrombocytosis phenotypes, but in most cases not both (Figure 3). This mutually exclusive red cell versus platelet phenotype was also found in recipients of *Ki* BM (Figure 3B), and therefore is not just a property of the *FF1* model. An interesting hypothesis for these observations is that individual HSCs may differ in lineage bias and expansion potential, which only becomes apparent when disease is initiated from a single HSC.

Engraftment and MPN was also observed in non-conditioned fully immunocompetent C57BL/6 mice when BM cells from *MxCre;Ki* mice that express mouse *Jak*-V617F were transplanted (Figure 4). In contrast, BM from *MxCre;FF1* mice failed to engraft in non-conditioned C57BL/6 mice even when five times higher number of BM were injected, presumably due to rejection directed against the human JAK2-V617F protein, despite 90% amino acid sequence identity with mouse JAK2 protein. We were able to reproduce the results obtained with *MxCre;Ki* also with *SclCre;Ki* BM transplanted into *SclCre* recipients, and even 50’000 BM cells per recipient were sufficient to engraft and initiate MPN. Thus, a small number of HSCs expressing the mouse *Jak*-V617F were able to initiate MPN in fully immunocompetent recipients. Using the “extreme limiting dilution analysis” algorithm,^23^ we estimated that 8’000 BM cells in transplantations into Rag2−/− recipients contained ∼1 functional LT-HSC, and thus these 50’000 BM cells are expected to contain ∼6 functional LT-HSCs per graft. The expansion of the Jak2-mutant HSC pool in the SclCre recipients was in the same range as observed in transplantations Rag2−/− recipients (Table 1).

Transplantations of wildtype BM into non-conditioned fully immunocompetent recipients are feasible when a large excess of BM cells is used.^27,28^ This led to the hypothesis that engraftment is limited by the number of HSC niches and large number of HSCs needed to be transplanted to compete for the few available empty niches.^29^ However, recent studies challenged this view and showed that HSCs that engrafted in non-conditioned recipients increased the total number of HSCs instead of replacing the numbers of endogenous resident HSCs, suggesting that numerous empty niches are available under steady state conditions.^30^ In our case, much smaller numbers of *Jak2*-mutant BM cells than wildtype BM cells were sufficient to engraft and expand in non-conditioned syngeneic hosts, since immune surveillance was bypassed when BM expressing the mouse *Jak2*-V617F was transplanted. In contrast, BM expressing the human *JAK2*-V617F was very effectively prevented from engrafting, suggesting T cell mediated rejection of the human JAK2 protein (Figure 5). The strong competitive advantage and the failure to be recognized by the immune surveillance may also explain why clonal hematopoiesis with *JAK2*-V617F in otherwise healthy individuals is surprisingly frequent, and with sensitive detection methods was reported to be ∼3%.^31^ Additional factors appear to limit the initial expansion of the *JAK2*-mutant clone, and contribute to keep the prevalence of *JAK2*-mutant CH higher than the prevalence of *JAK2*-mutant MPN. Nestin-positive MSCs appear to exert a negative effect,^32^ whereas inflammation is favoring the expansion of *JAK2*-V617F HSCs.^33,34^ Our non-conditioned model will now allow to study these aspect in the absence of irradiation and the following cytokine storm. Failure of mouse JAK2-V617F protein to be recognized by the mouse immune system also suggest that immuno-therapeutic approaches against *JAK2*-mutant MPN will be more challenging than immuno-therapies against driver proteins, such as CALRdel52, that represent neo-antigens.

Our data indicate that the *Jak2*-V617F mutation provides a strong competitive advantage to the HSCs, provided that the mutated cells are not eliminated by the immune surveillance, as was the case with the human *JAK2*-V617F. Transplantations into non-conditioned hosts at limiting dilutions of HSCs opens new ways for studying *JAK2*-mutant clonal expansion and disease initiation avoiding damage to the microenvironment by irradiation and increased cytokine levels due to post radiation aplasia.

## AUTHOR CONTRIBUTIONS (CRediT Classification)

Quentin Kimmerlin: Conceptualization, Investigation, Methodology, Visualization, Writing – Original Draft Preparation and Writing – Review and Editing.

Morgane Hilpert: Conceptualization, Investigation, Methodology and Visualization.

Nils Hansen: Conceptualization, Investigation and Methodology.

Alexandre Guy: Conceptualization, Investigation and Methodology.

Marc Usart: Conceptualization, Investigation and Methodology.

Jan Stetka: Conceptualization, Investigation and Methodology.

Patryk Sobieralski: Investigation and Methodology.

Tiago Almeida Fonseca: Investigation and Methodology.

Hui Hao-Shen: Investigation and Methodology.

Radek C. Skoda: Conceptualization, Funding Acquisition, Project Administration, Resources, Supervision, Visualization, Writing – Original Draft Preparation and Writing – Review and Editing.

## DATA SHARING AND AVAILABILITY

For original data and reagents, please contact.

## ACKNOWLEDGMENTS

We thank the members of the laboratory for helpful discussions and critical reading of our manuscript.

