## Supplemental Figure S1 for "Single *JAK2*-V617F hematopoietic stem cells can initiate MPN in transplantations into non-conditioned recipient mice"

### Supplemental Figure 1 (related to Figure 1)

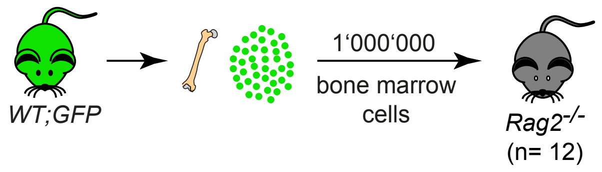


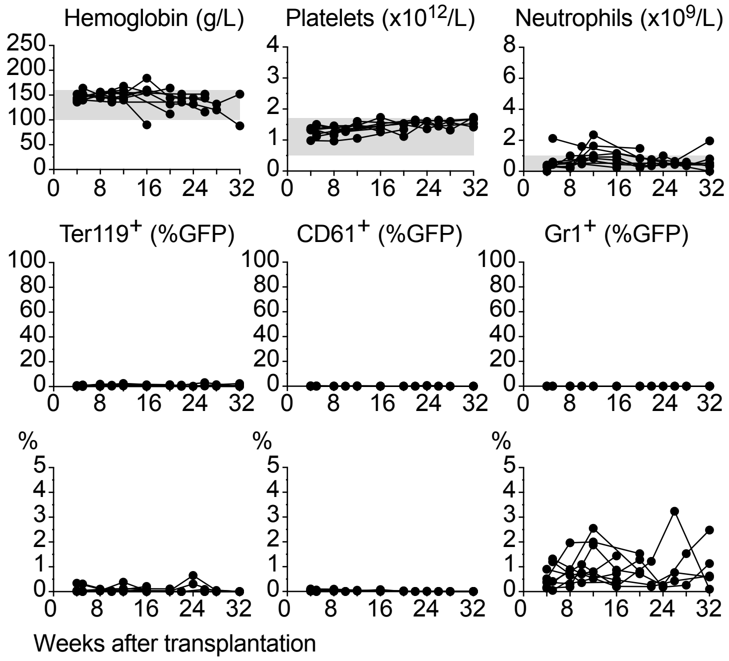


**Legend to Supplemental Figure S1.**

A total of 1'000'000 unfractionated bone marrow cells obtained from a wildtype donor (*WT;GFP)* were transplanted into non-conditioned *Rag2$^{-/-}$* recipients. (**A**) Schematic drawing of the experimental setup. (**B**) Blood counts and GFP chimerism of non-conditioned *Rag2$^{-/-}$* recipients, represented for each individual mouse. Chimerism is also plotted in an expanded range below to visualize low degree of engraftment.
