## Supplemental Figure S2 for "Single *JAK2*-V617F hematopoietic stem cells can initiate MPN in transplantations into non-conditioned recipient mice"

### Supplemental Figure 2 (related to Figure 1)

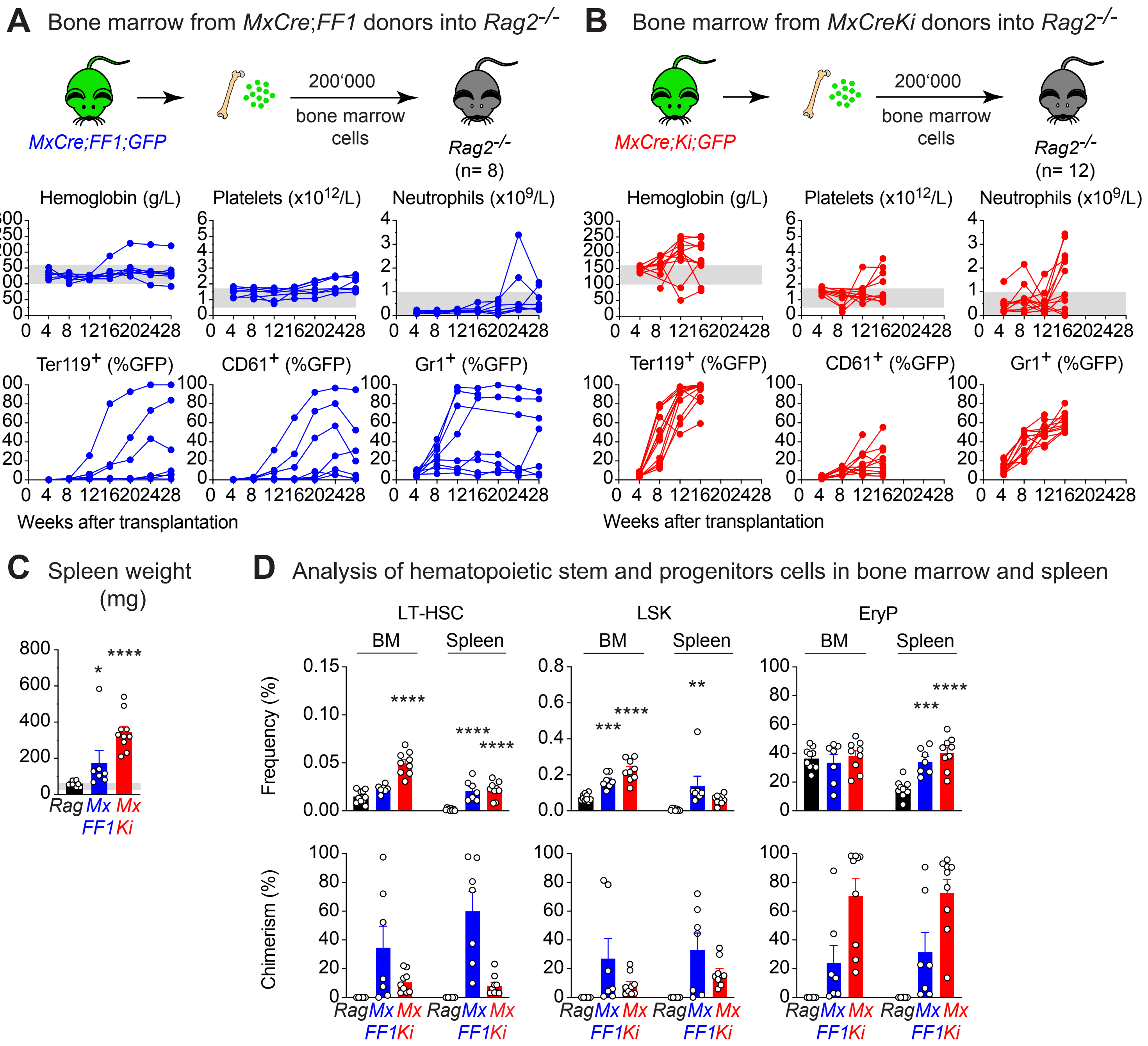


**Legend to Supplemental Figure S2.**
(**A and B**) A total of 200’000 unfractionated bone marrow (BM) cells obtained from *MxCre;FF1;GFP* (blue) or *MxCre;Ki;GFP* (red) donors were transplanted into non-conditioned *Rag2*$^{-/-}$ recipients. Experimental design and time course of blood counts and GFP chimerism. Each line represents an individual mouse. (**C**) Spleen weight of *Rag2*$^{-/-}$ recipients at terminal analysis at 28 weeks and 16 weeks, respectively. Non-transplanted *Rag2*$^{-/-}$ mice (black) were plotted for comparison. (**D**) Frequencies and GFP chimerism of LT-HSCs (lin- ckit+ Sca1+ CD150+ CD48-), LSKs (lin- ckit+ Sca1+) and EryP (erythroid progenitors, CD71+ Ter119+). Non-transplanted *Rag2*$^{-/-}$ mice (black) were plotted for comparison. Statistical analyses were performed using one-way ANOVA followed by Fisher’s LSD test, with the non-transplanted *Rag2*$^{-/-}$ mice serving as the reference for comparisons. *P < .05; **P < .01; ***P < .001; ****P < .0001.
