## Supplementary figures and images for "Single *JAK2*-V617F hematopoietic stem cells can initiate MPN in transplantations into non-conditioned recipient mice"

### Supplemental Figure S3

#### Supplemental Figure 3 (related to Figure 2)

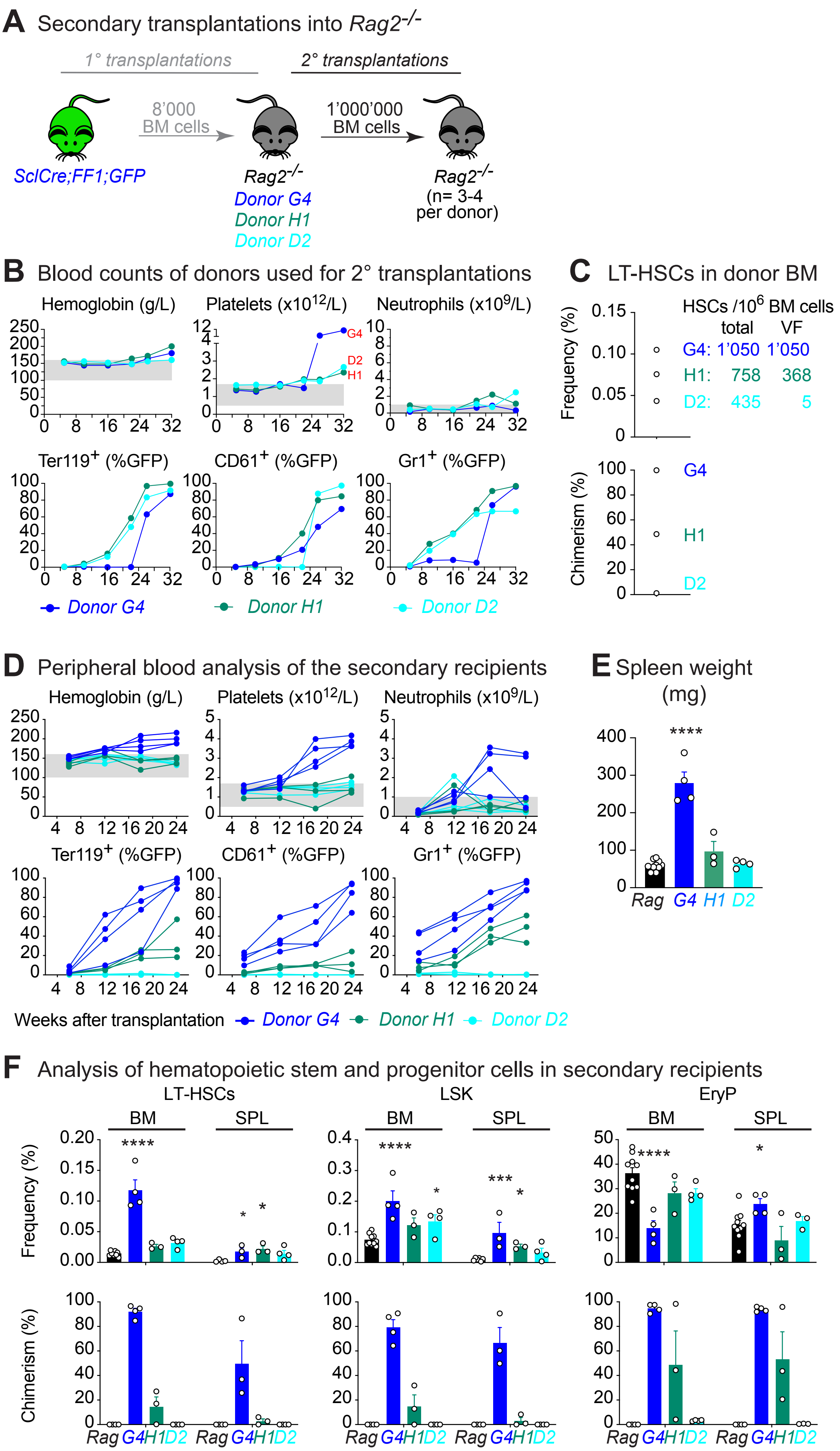
