## Supplemental Figure S3 (Figure Legend) for "Single *JAK2*-V617F hematopoietic stem cells can initiate MPN in transplantations into non-conditioned recipient mice"

**Legend to Supplemental Figure S3.**

(**A**) Experimental design of secondary transplantations. (**B**) Blood counts of the three non-conditioned *Rag2*$^{-/-}$ recipients transplanted with 8’000 bone marrow (BM) cells from *SclCre;FF1;GFP* that were used as donor mice for secondary transplantations. (**C**) Frequency and GFP chimerism of LT-HSCs (lin- ckit+ Sca1+ CD150+ CD48-) in the three donors that were used for secondary transplantations. (**D**) Time course of blood counts and GFP chimerism of secondary recipients. Each line represents one individual mouse. (**E**) Spleen weight of non-conditioned *Rag2*$^{-/-}$ secondary recipients at terminal analysis. Non-transplanted *Rag2*$^{-/-}$ mice (black) were plotted for comparison. (**F**) Frequencies and GFP chimerism of LT-HSCs, LSKs (lin- ckit+ Sca1+) and EryP (erythroid progenitors, CD71+ Ter119+) in secondary recipients. Non-transplanted *Rag2*$^{-/-}$ mice (black) were plotted for comparison. Statistical analyses were performed using one-way ANOVA followed by Fisher’s LSD test, with the non-transplanted *Rag2*$^{-/-}$ mice serving as the reference for comparisons. *P < .05; **P < .01; ***P < .001; ****P < .0001.
